# Potential Hematotoxicity and Genotoxicity of Multi-Herbal Formulations in Albino Mice (*Mus musculus*)

**DOI:** 10.1101/2020.07.31.231407

**Authors:** S. Kehinde, S. M. Adebayo, A. L. Adesiyan, E. A. Kade, K. Gurpreet

**Affiliations:** Department of Cell biology and Genetics, University of Lagos (UNILAG), Akoka, Lagos, Nigeria; Department of Microbiology, University of Lagos (UNILAG), Akoka, Lagos, Nigeria; Department of Health and Health Care Administration, Swami Rama Himalayan University (SRHU), Dehradun, Indian

**Keywords:** Multi-herbal Formulations, Genotoxicity, Hematotoxicity, DNA fragmentation

## Abstract

The current increase in the use of multi-herbal remedies coupled with loose regulation on public access to these products underscore research efforts to evaluate their biochemical effect, noting that many of the herbal medicines lack scientific evidence to support their medicinal claims. Objective: We therefore investigated the potential genotoxicity and hematotoxicity of commonly consumed multi-herbal formulations (YoyoBitters™, Ogidiga™ and BabyOku™) in Lagos, Nigeria, in experimental mice. Methods: Fifty (50) adult female albino mice were randomly selected and distributed into 5 groups of 10 mice each. Two mL/kg body weight of distilled water were orally administered to the control groups while BabyOku™, YoyoBitters™ and Ogidiga™ herbal formulations were administered to the experimental groups at doses of 2 mL/kg body weights. Results: A dose- and tissue-dependent increase in induction of apoptotic DNA fragmentation was observed in the triherbal groups relative to control groups. Also, an increase in micronucleated polychromatic erythrocytes was formed in a dose-dependent manner in the multi-herbal groups when compared with the control groups. Conclusion: From our findings, multi-herbal formulations may possess hematotoxic and genotoxic potentials in mice.

## 1. Introduction

Medicinal plants are a major source of active drugs from nature. The use of plant parts in treating diseases is universal, it is often more affordable and believed to be effective than the conventional drugs. Most of these medicinal plants are eaten or used for their rich phytochemical constituents, which provide both preventive and curative properties to consumers against various diseases [1]. In recent years, there has been increase in the popularity enjoyed by herbal remedy usually prepared by mixing various medicinal plant species [2].

In Nigeria, the last few years have witnessed an increase in the demand of herbal remedies. In spite of the wide patronage enjoyed by herbal remedies, little or no empirical data exist to support medicinal claims. Also, there are no scientific data on safety and toxicity profiles of these herbals [2-4]. Herbal remedy mixtures such as YoyoBitters™, Swedish Bitters™, Fijk™, OsomoBitters™, Alomo™, Oroki™ among others have become a common sight in many Nigerian homes. All of these herbals have highly praised medicinal benefits but only few, if any, have empirical data to support medicinal claims. However, recent studies have demonstrated the need to subject some of the herbal mixtures to scientific examination, at least in part to ascertain safety limits [2-6], more so government regulation of herbal medicine is not so stringent when compared to conventional drugs. Furthermore, microbial contaminants and higher level of heavy metals which could be detrimental to human health have been demonstrated in several herbal remedies [4, 7]. Toxicity may be relatively unknown even for efficient and documented herbal medicinal products; in fact, unlike conventional drug research and development, the toxicity of traditional herbal medicinal products is not often evaluated. [8-10]. However, the majority of the population does not pay attention to its toxicity, believing that if these products have been used for years, they should be devoid of toxic substance. [11-14]. All of these factors serve to encourage the imperativeness for empirical data on either the safety or toxicity margin of herbal mixtures being marketed and promoted to the Nigerian populace.

Ogidiga™ and BabyOku™ herbal mixtures are very popular among the Nigerian populace. The constituents of Ogidiga™ according to its label include ethanol, water, sugar, lemon, garlic, ginger and *combretaceae*. [4]. BabyOku™ contains carene, a medicinal component of herbal drugs, eugenol (alcohol), fatty acids such as propenoic acid and nonanoic acid and cyclohexanemethanol. Other active ingredients in it are water, ethanol, caramel, herbal flavour, extracts such as angelia root, *Cassia sanna* (sic) leaf, rhubarb root and aloe. Their bitterness are claimed to boost libido, cure pile, malaria, clear toxins among others, but there are no scientific data on any of these herbals to support these medicinal claims or otherwise [4]. There are increasing reports that several plants contain toxic, genotoxic and carcinogenic compounds [15-17]. Chromosomal aberrations in plants and animals are hallmarks of genome instability which may lead to genetic related diseases and congenital abnormalities [18, 19]. This study therefore investigated in rats, the potential genotoxicity and hematotoxicity, of two commonly consumed multi-herbal formulations (Ogidiga™ and Baby Oku™) in Lagos, Nigeria.

## 2. Materials and Methods

### 2.1. Experimental Animals

Fifty (50) adult female mice weighing 100-150g, used for this study, were purchased at the Nigeria Institute for Medical Research (NIMR), Yaba, Lagos. They were acclimatized for two weeks before the onset of the experiment. The animals were housed in wooden cages with good aeration, in a room with average illumination with 12:12-hour light:dark cycle and they were given free access to water and supplied with standard pellet *ad-libitum*.

### 2.2. Test Substances/Formulations

The Multi-herbal formulations were purchased at a liquor store at Bariga Area, Akoka, Nigeria. The formulations used for the experiment were Fanta® (a non–alcoholic non-multiherbal formulation), YoyoBitters™ (a multi-herbal non-alcoholic formulation), BabyOku™ and Ogidiga™ (a multi-herbal alcoholic formulations).

### 2.3. Experimental Design

The rats were randomly selected and assigned into 5 groups based on the type and amount of formulation/test substance administered. Each group contained 10 mice each based on the duration of administration. Two (2) mL/kg body weight of distilled water, Fanta®, and YoyoBitters™ were orally administered to the control groups while BabyOku™ and Ogidiga™ polyherbal formulations were administered to the experimental groups at doses of 2 mL/kg and 3 mL/kg body weights. The experimental mice were sacrificed at intervals of day 0, day 8, day 16, day 24 and day 32 of the administration of the test substances by cervical dislocation. Two (2) hours prior to sacrifice, each rat was injected with colchicine (prepared in distilled water) at a dose of 1 mL/100 g body weight intraperitoneally, for mitotic arrest. Mice were dissected and blood samples were collected with heparinized syringes via the abdominal artery and immediately transferred to heparinized tube and kept on ice for full and differential blood counts. Tissues (liver, kidneys, heart, brain, lungs, ovaries, uterus, spleen) were harvested, washed in ice-cold normal saline and stored at −20 °C for genotoxicity experiments. Femur from both legs were quickly harvested and immediately used for the micronucleus assay.

### 2.4. Full and Differential Blood Counts

Samples of EDTA-anticoagulated blood were collected and stored in a cool box, at approximately 4 °C, and delivered to a local processing field laboratory within two hours of collection. Full and differential blood counts were analyzed on a Coulter LH700 series Hematology analyzer (Beckman Coulter, Miami, USA).

### 2.5. Micronucleus Assay

The femurs from each of the animals were removed and bone marrow was aspirated with a syringe and microscopic slides prepared according to Matter and Schmid [28]. The slides were then fixed in absolute methanol (BDH Chemical Ltd, Poole, England), air-dried, pretreated with May-Grunwald solution (Sigma-Aldrich, procedure No GS-10) and air-dried. The dried slides were stained in 5% Giemsa solution, and immersed in phosphate buffer 0.01 mol L-1 (pH 6.8) for 30 s. Thereafter, they were rinsed in distilled water, air-dried, and mounted. The slides were scored at x 100 magnification under a Nikon E200 light microscope (Opto-Edu Co., Ltd, Beijing, China) for micronucleated polychromatic erythrocytes (mPCEs).

### 2.6. DNA Fragmentation Assay

The method of Wu et al. [29] was used. The tissues were homogenized in 10 volumes of a lysis buffer (pH 8.0) consisting of 5 *mM* Tris-HCI, 20 *mM* EDTA and 0.5% (w/v) t-octylphenoxypolyethoxyethanol (Triton X-100). 1 mL aliquots of each sample were centrifuged at 27,000 *g* for 20 minutes to separate the intact chromatin (pellet) from the fragmented DNA (supernatant). The supernatant was decanted and saved, and the pellet was resuspended in 1 mL of Tris buffer (pH 8.0) consisting of 10 *mM* Tris-HCI and 1 *mM* EDTA. The pellet and supernatant fractions were assayed for DNA content using a diphenylamine reaction. Optical density was read at 620 nm with spectrophotometer. The results were expressed as a percentage of fragmented DNA divided by total DNA.

### 2.7. Statistical Analysis

Data were analyzed by one-way analysis of variance (ANOVA), followed by Duncan Multiple Range Test to test for significant differences among the groups of rats using SPSS 16.0. Data were expressed as mean ± standard error of mean. P values less than 0.05 were considered statistically significant.

### 3. Results

### 3.1. Alterations in Haematological Parameters in multi-herbal Formulations Treated mice

There was a significant decrease in haematological parameters (Table 1, 2, 3 and 4) by all the multi-herbal formulations compared with the control (DH2O) group (p□0.05). A statistical dose dependent decrease in the hemoglobin concentration and percentage packed cell volume; red blood cell, white blood cell, lymphocytes, neutrophil and platelets counts were observed (2 mL/kg bodyweight) among rats treated with BabyOku™ and Ogidiga™ multi-herbal formulations (p□0.05). It was observed that the degree of alteration produced by the polyherbal formulations is Ogidiga™ □ BabyOku™. There was a significant difference in the degree of alterations in haematological parameters induced by the multi-herbal formulations relative to Fanta®, Yoyo bitters™ or treated groups (p□0.05).

**Table 1.** Hemoglobin concentration, red blood cells and platelets counts in multi-herbal formulations treated mice.

|  | HEMOGLOBIN CONCENTRATION (g/dl) |  |  | RED BLOOD CELLS (x 1012 g/L) |  |  | PLATELETS COUNTS (x103 cells/L) |  |  |
| --- | --- | --- | --- | --- | --- | --- | --- | --- | --- |
|  | DAY 0 | DAY 16 | DAY 32 | DAY 0 | DAY 16 | DAY 32 | DAY 0 | DAY 16 | DAY 32 |
| DH20 | 16.36±0.15 ab | 16.78±0.08 a | 16.74±0.15 a | 7.63±0.01 bc | 7.37±0.02 b | 7.07±0.02 b | 803.00±2.23 d | 860.60±1.95 a | 851.80±1.79a |
| Fanta® | 16.26±0.1<br>1 ab | 16.38±0.31<br>b | 16.26±0.11<br>b | 7.86±0.0<br>1 a | 7.47±0.01<br>a | 7.90±0.02<br>a | 756.80±1.92<br>e | 853.80±1.48<br>b | 811.40±2.30b |
| Yoyo<br>bitters<br>™ | 16.36±0.1<br>1 ab | 15.26±0.11<br>c | 14.26±0.18<br>c | .55±0.01<br>cd | 6.73±0.05<br>c | 5.67±0.02<br>c | 904.80±1.48<br>a | 789.80±1.30<br>c | 767.20±41.69<br>c |
| BO2 | 16.30±0.1<br>5 ab | 11.70±0.16<br>e | 4.26±0.11f | 7.69±0.0<br>1 b | 5.81±0.06<br>e | 2.55±0.04<br>e | 912.40±1.81<br>a | 604.80±2.05<br>f | 393.60±1.14e |
| OG2 | 16.48±0.2<br>1 ab | 10.30±0.16<br>g | 3.26±0.15h | 7.93±0.0<br>1 a | 5.32±0.01<br>h | 2.43±0.02<br>f | 756.80±1.92<br>e | 403.80±2.28<br>j | 212.40±2.07h |

Values are expressed as mean ± standard deviation (n=5); values bearing different alphabets are significantly different while those with similar alphabets are insignificantly different ( $p < 0.05$ ). BO2= 2 mL/kg body weight of BabyOku™, OG2= 2 mL/kg body weight of Ogidiga™

**Figure 1:**
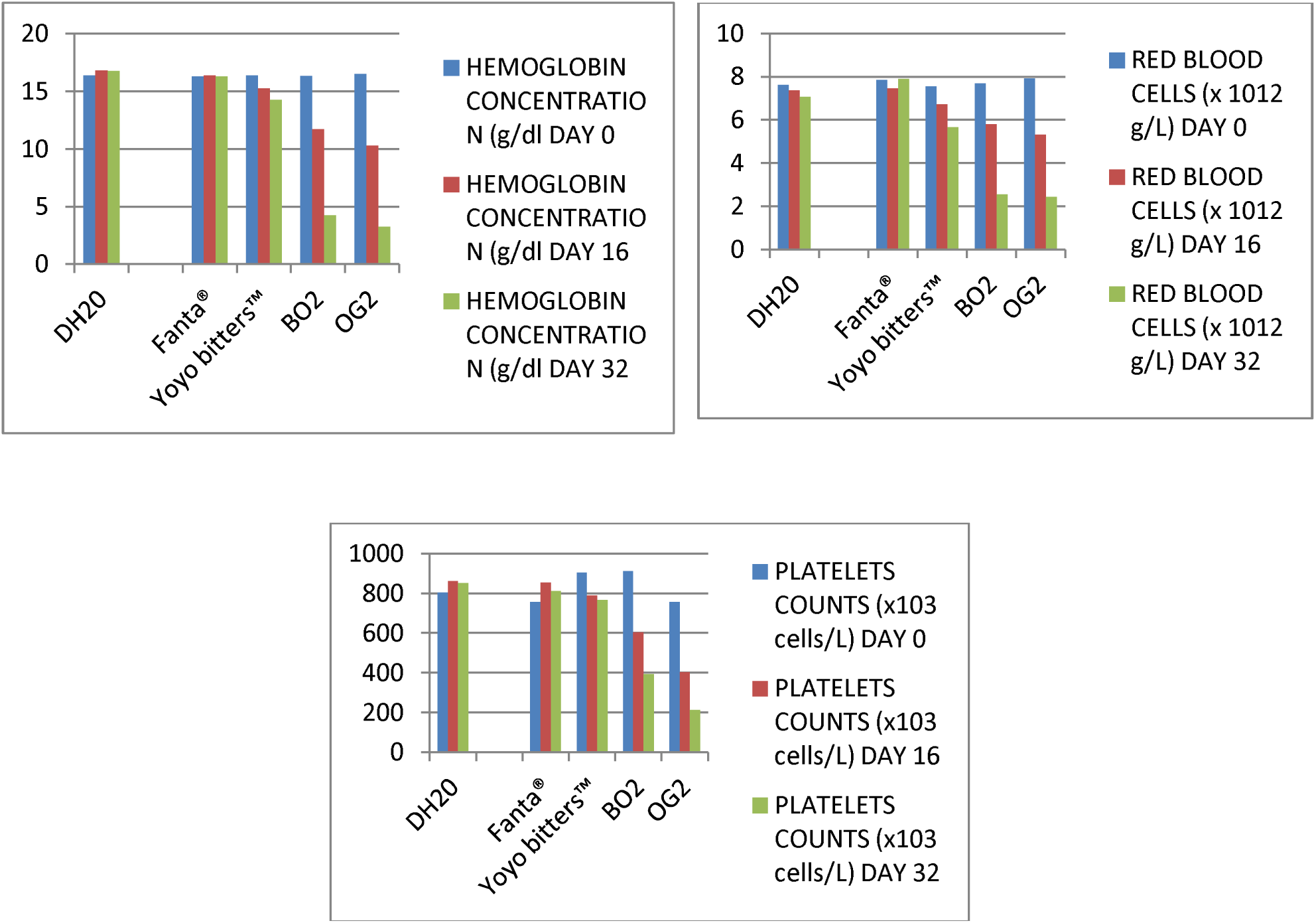
Hemoglobin concentration, red blood cells and platelets counts in multi-herbal formulations treated mice.

**Table 2.**
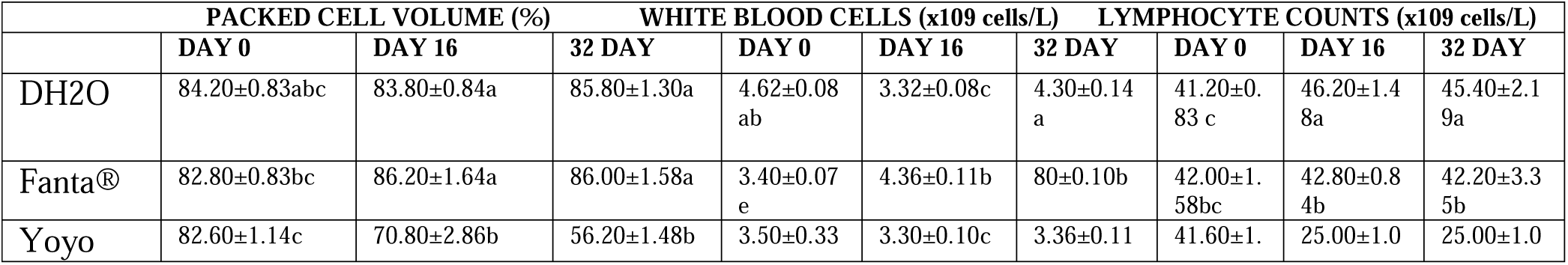
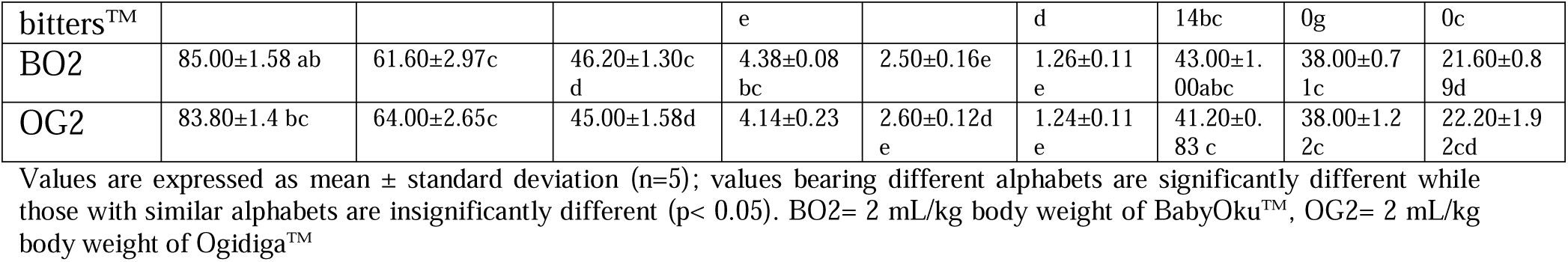
Packed cell volume, white blood cells and lymphocyte counts in herbal formulations treated mice.

**Table 3.**
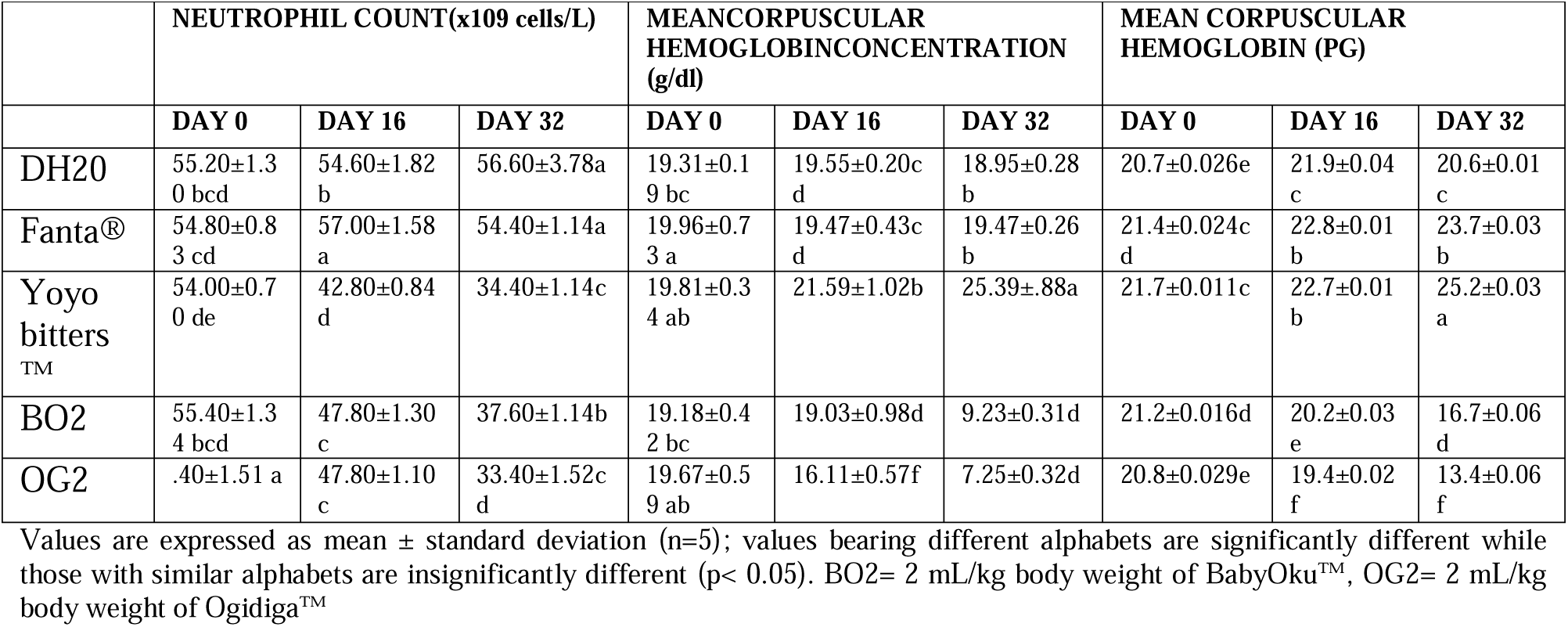
Neutrophil count, mean corpuscular hemoglobin concentration (MCHC) and mean corpuscular hemoglobin (MCH) in multi-herbal formulations treated mice.

**Table 4.**
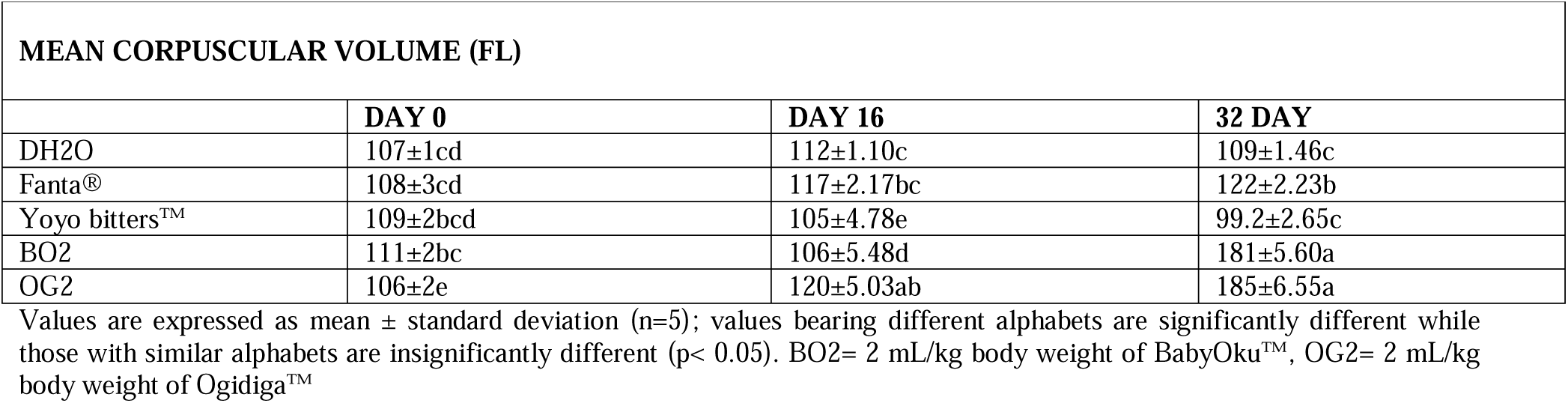
Alterations in mean corpuscular volume (MCV) in herbal formulations treated mice.

| MEAN CORPUSCULAR VOLUME (FL) |  |  |  |
| --- | --- | --- | --- |
|  | DAY 0 | DAY 16 | 32 DAY |
| DH20 | 107±1cd | 112±1.10c | 109±1.46c |
| Fanta® | 108±3cd | 117±2.17bc | 122±2.23b |
| Yoyo bitters™ | 109±2bcd | 105±4.78e | 99.2±2.65c |
| BO2 | 111±2bc | 106±5.48d | 181±5.60a |
| OG2 | 106±2e | 120±5.03ab | 185±6.55a |

Values are expressed as mean ± standard deviation (n=5); values bearing different alphabets are significantly different while those with similar alphabets are insignificantly different (p< 0.05). BO2= 2 mL/kg body weight of BabyOku™, OG2= 2 mL/kg body weight of Ogidiga™

### 3.2. Genotoxic Potentials of Baby Oku™ and Ogidiga™ on Experimental Mice

There was a significant increase in induction of DNA fragmentation (Tables 5 to 9) by all the herbal formulations compared with the control (DH2O) group (p□0.05). A statistical dose dependent increase in induction of DNA fragmentation were observed (2 mL/kg body weight) among mice treated with BabyOku™ and Ogidiga™ multi-herbal formulations (p□0.05). It was observed that the degree of induction of apoptotic DNA fragmentation by the herbal formulations is in the following order: BabyOku™ □ Ogidiga™. There was a significant increase in the degree of induction of DNA fragmentation by the herbal formulations relative to Fanta®, Yoyo bitters™ or treated groups (p□0.05). A varying degree of percentage DNA fragments were observed for each herbal formulation. The trend for BabyOku™ was heart > kidney > liver > brain >lungs, Ogidiga™ had heart > kidney >liver > brain > lungs.

**Table 5.**
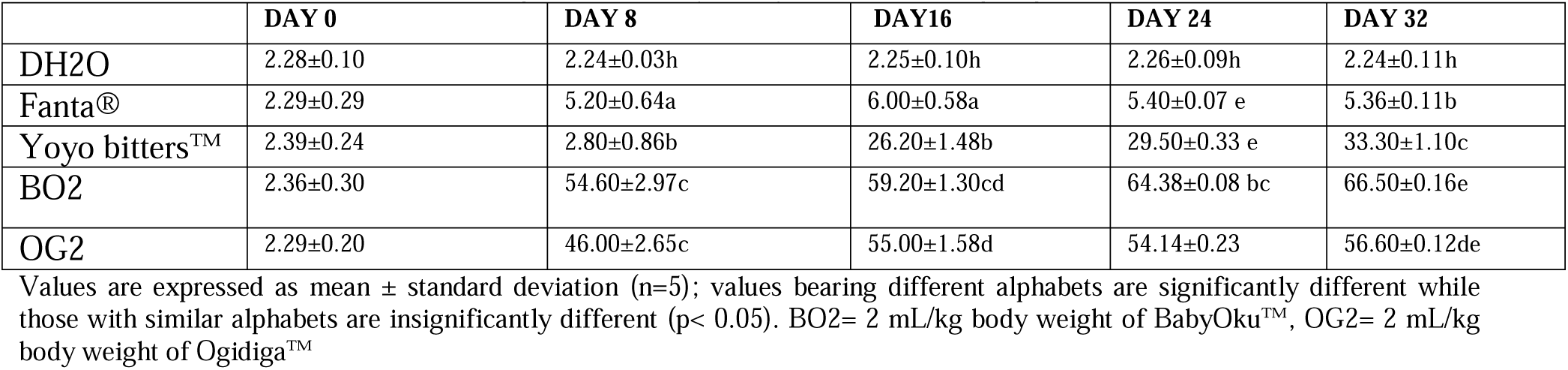
Induction of brain DNA fragmentation by BabyOku™ and Ogidiga™ in experimental mice.

**Table 6.** Induction of hepatic DNA fragmentation by BabyOku™ and Ogidiga™ in experimental mice.

|  | DAY 0 | DAY 16 | 32 DAY | DAY 0 | DAY 16 |
| --- | --- | --- | --- | --- | --- |
| DH2O | 2.28±0.10 | 2.24±0.03h | 85.80±1.30a | 4.62±0.08 ab | 3.32±0.08c |
| Fanta® | 2.29±0.29 | 5.20±0.64a | 86.00±1.58a | 33.40±0.07 e | 34.36±0.11b |
| Yoyo bitters™ | 2.39±0.24 | 21.80±0.86b | 56.20±1.48b | 53.50±0.33 e | 63.30±0.10c |
| BO2 | 2.36±0.30 | 39.60±2.97c | 46.20±1.30cd | 64.38±0.08 bc | 72.50±0.16e |
| OG2 | 2.29±0.20 | 34.00±2.65c | 45.00±1.58d | 47.14±0.23 | 52.60±0.12de |

Values are expressed as mean ± standard deviation (n=5); values bearing different alphabets are significantly different while those with similar alphabets are insignificantly different ( $p < 0.05$ ). BO2= 2 mL/kg body weight of BabyOku™, OG2= 2 mL/kg body weight of Ogidiga™

**Table 7.** Induction of lungs DNA fragmentation by BabyOku™ and Ogidiga™ in experimental mice.

|  | DAY 0 | DAY 16 | 32 DAY | DAY 0 | DAY 16 |
| --- | --- | --- | --- | --- | --- |
| DH2O | 1.20±0.83abc | 1.80±0.84a | 1.80±1.30a | 1.62±0.08 ab | 1.32±0.08c |
| Fanta® | 1.80±0.83bc | 2.20±0.64a | 2.00±0.58a | 2.40±0.07 e | 2.36±0.11b |
| Yoyo bitters™ | 1.60±0.14c | 10.80±2.86b | 12.20±1.48b | 23.50±0.33 e | 40.30±0.10c |
| BO2 | 2.00±0.58 ab | 20.60±2.97c | 26.20±1.30cd | 30.38±0.08 bc | 82.50±0.16e |
| OG2 | 2.80±0.4 bc | 14.00±2.65c | 19.00±1.58d | 24.14±0.23 | 32.60±0.12de |

Values are expressed as mean ± standard deviation (n=5); values bearing different alphabets are significantly different while those with similar alphabets are insignificantly different ( $p < 0.05$ ). BO2= 2 mL/kg body weight of BabyOku™, OG2= 2 mL/kg body weight of Ogidiga™

**Table 8.**
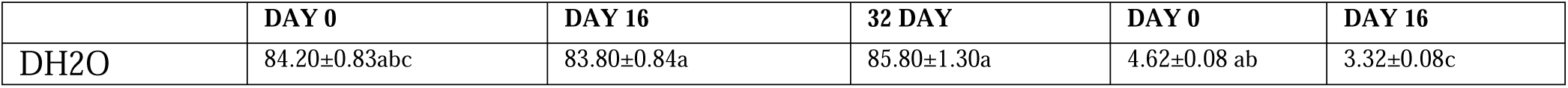
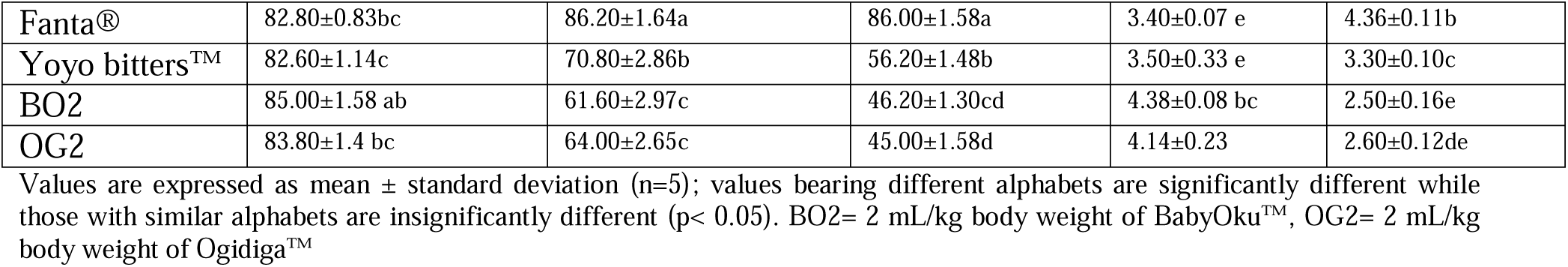
Induction of cardiac DNA fragmentation by BabyOku™ and Ogidiga™ experimental mice.

|  | DAY 0 | DAY 16 | 32 DAY | DAY 0 | DAY 16 |
| --- | --- | --- | --- | --- | --- |
| DH2O | 84.20±0.83abc | 83.80±0.84a | 85.80±1.30a | 4.62±0.08 ab | 3.32±0.08c |
| Fanta® | 82.80±0.83bc | 86.20±1.64a | 86.00±1.58a | 3.40±0.07 e | 4.36±0.11b |
| Yoyo bitters™ | 82.60±1.14c | 70.80±2.86b | 56.20±1.48b | 3.50±0.33 e | 3.30±0.10c |
| BO2 | 85.00±1.58 ab | 61.60±2.97c | 46.20±1.30cd | 4.38±0.08 bc | 2.50±0.16e |
| OG2 | 83.80±1.4 bc | 64.00±2.65c | 45.00±1.58d | 4.14±0.23 | 2.60±0.12de |

Values are expressed as mean ± standard deviation (n=5); values bearing different alphabets are significantly different while those with similar alphabets are insignificantly different (p< 0.05). BO2= 2 mL/kg body weight of BabyOku™, OG2= 2 mL/kg body weight of Ogidiga™

**Table 9.** Induction of renal DNA fragmentation by BabyOku™ and Ogidiga™ in experimental mice.

|  | DAY 0 | DAY 8 | DAY16 | DAY 24 | DAY 32 |
| --- | --- | --- | --- | --- | --- |
| DH2O | 2.28±0.10 | 2.24±0.03h | 2.25±0.10h | 2.26±0.09h | 2.24±0.11h |
| Fanta® | 2.29±0.29 | 5.20±0.64a | 6.00±0.58a | 5.40±0.07 e | 5.36±0.11b |
| Yoyo bitters™ | 2.39±0.24 | 2.80±0.86b | 26.20±1.48b | 29.50±0.33 e | 33.30±1.10c |
| BO2 | 2.36±0.30 | 54.60±2.97c | 59.20±1.30cd | 64.38±0.08 bc | 66.50±0.16e |
| OG2 | 2.29±0.20 | 46.00±2.65c | 55.00±1.58d | 54.14±0.23 | 56.60±0.12de |

Values are expressed as mean ± standard deviation (n=5); values bearing different alphabets are significantly different while those with similar alphabets are insignificantly different (p< 0.05). BO2= 2 mL/kg body weight of BabyOku™, OG2= 2 mL/kg body weight of Ogidiga™

## 4. Discussion

Medicinal plants have been widely used by both ancient and modern man of all cultures for treating different ailments. A single plant processed in different formulations can be used to cure a wide range of diseases [30]. The use of herbal materials in alternative medicine plays important roles in primary health care for most African countries, mainly due to their culture and beliefs. Despite the profound therapeutic advantages presented by many of these medicinal plants, some still exhibit some systemic toxicity, genotoxicity and carcinogenicity potentials [17, 33-35]. Therefore, there is the need for more information on the toxicological profile of many of the herbal supplements used in the complementary and alternative medicine in Nigeria and most other countries of the world [19]. However, the historic role of medicinal herbs in the treatment and prevention of diseases and in the development of pharmacology do not assume their safety for uncontrolled use by an uninformed public [31, 32]. This study presents the hematological alterations and genotoxic effects induced by the two commonly consumed multi-herbal formulations (MHFs) (Ogidiga™ and BabyOku™) in lagos, Nigeria on experimental mice. Clinical signs of toxicity observed mainly due to the administration of Multi-herbal formulations in mice are systemic toxicity.

Hematological testing in rodents during toxicity and safety evaluation is generally acknowledged as integral part of systemic toxicity assessment [36]. Significant decrease in packed cell volume; hemoglobin, white blood cells, lymphocytes, neutrophils and platelet counts in Ogidiga™ and BabyOku™ exposed mice showed that the multi-herbal formulations induced marked hematotoxic effects in the rodents. Alterations in hematological biomarkers suggest that the component phytochemicals of the various multi-herbal formulations affected hematopoiesis in the bone marrow system of the MHFs exposed mice [33, 34, 37].

Anemia is a reduction in the number of erythrocytes, hemoglobin, or both, in the circulating blood. It resulted from excessive red blood cell (RBC) destruction, RBC loss, or decreased RBC production and is a manifestation of an underlying disease process. Therefore, the response to treatment of anemia is transient unless the underlying disease process is addressed [38]. Although it was stated that toxic plants do not produce a direct effect on white blood cells, such as neutrophils, lymphocytes, eosinophils, and monocytes [39], the results of this study showed otherwise. Excessive consumption of a wide variety of plants or their products has been found to cause hypo-proliferative or nonregenerative anemia, which is a stem cell disorder characterized by reduced bone marrow production of all blood components in the absence of a primary disease process infiltrating the bone marrow or suppressing hematopoiesis [40]. It shows that continuous consumption of these formulations may produce these effects in animals. It may also mean that the principal function of white blood cells, which is to defend against invading organisms, will be compromised [38, 39, 41]. Since one of the pathways leading to apoptosis involves DNA degradation, it is worth stating with emphasis that multi-herbal formulations might trigger apoptosis by damaging genetic material.

## 5. Conclusions

We therefore concluded that Ogidiga™ and BabyOku™, the commonly consumed multi-herbal formulations in Lagos, Nigeria has strong potential to induce genotoxicity and hematotoxicity in experimental mice as evident from increased DNA fragmentation, induction of micro-nucleated polychromatic erythrocytes (mPCEs) and reduction in hematological biomarkers.

